# Identifying Therapeutic Strategies in IgA Nephropathy through Comprehensive Transcriptomic Characterization

**DOI:** 10.1101/2021.08.18.456879

**Authors:** Kenneth Ofori, Amin Yakubu, Alex J. Rai

## Abstract

IgA nephropathy (IgAN) is an autoimmune disease and the most common primary glomerulonephritis. The four-hit hypothesis describes mechanism of the disease, from synthesis of galactose deficient IgA (GD-IgA), to recognition of GD-IgA by anti-glycan antibodies and deposition of the formed immune complex in the mesangium. Complement and coagulation cascade activation ensues, resulting in mesangial activation and cytokine release, podocyte injury, mesangial sclerosis and tubulointerstitial damage. Currently, there is no disease cure, and 30-40% of patients progress to end stage renal disease.

Using complementary bioinformatic approaches, we demonstrate different levels of deviation of the transcriptome of the glomerulus in IgAN from normal, with the aim of identifying therapeutic targets. Approaches used herein include, deconvolution of the transcriptome to estimate immune constitution, co-regulation-based functional analysis of differentially expressed genes, modular co-expression analysis, network analysis of metabolic pathways and differential gene correlation analysis.

We describe the immune composition in IgAN and the relatively low fold changes of the abundance of different immune cells and strength of immune signatures compared with control. Additionally, we identify enrichment of the intestinal network for IgA synthesis, repression of expression and dysregulation of networks of amino acid metabolism and PPAR signaling pathways in IgAN glomeruli. We also find loss of correlation between expression of matrix synthesizing and matrix degrading genes in IgAN.

We conclude by discussing how therapies based on some nodes in these altered pathways described have been shown to be efficacious in IgAN and/or other inflammatory diseases and the potential of others in effective treatment.

## Introduction

Immunoglobulin A nephropathy (IgAN), also known as Berger’s disease, is the most common primary glomerulonephritis worldwide.^1^ The disease presents as a nephritic syndrome, consisting of hematuria, hypertension, albuminuria and elevation of serum creatinine. Significant progress has been made in elucidating the etiology of IgAN. The four-hit hypothesis explains the sequence of events from aberrant IgA synthesis to kidney disease. The first hit in IgAN is abnormal O-glycosylation of the hinge region of IgA heavy chain resulting in galactose deficient IgA 1(GD-IgA1). Absence of galactose is thought to expose normally sequestered terminal and sialylated N-acetylgalactosamine (GalNAc) residues on the hinge region, leading to their detection by naturally occurring anti-glycan IgG or IgA autoantibodies.^2–4^ Binding of these autoantibodies to GD-IgA forms immune complexes which are subsequently deposited in the mesangium of the kidney.^1^ The deposited immune complex fixes complement and attracts leucocytes with cytokine production, causing mesangial activation, proliferation and extracellular matrix deposition.^5,6^ Podocyte injury and loss, and tubulointerstitial inflammation and damage result from the elaborated cytokines from activated mesangium.^7,8^ The histologic picture consists of mesangial proliferation, endocapillary proliferation, glomerulosclerosis, tubular atrophy and interstitial fibrosis, and crescent formation in severe cases.^9^ In spite of the progress made in understanding its etiology, there is currently no disease specific therapy for IgAN. About 30-40% of patients slowly develop end stage renal disease in 20-30 years.^1^ There is therefore a need to identify other mechanistic features of the disease that can be targeted by therapy.

Analysis of the transcriptome has contributed considerably in elucidating disease mechanisms and progression. A common approach in analyzing the transcriptome of disease is by differential gene expression, comparing disease with normal to identify aberrantly expressed genes.^10,11^ While differential gene expression has been useful in studying disease including IgAN, it has been shown that transcriptomic dysregulation in disease involves a concerted alteration of groups of co-expressed and co-regulated genes, and therefore network approaches provide more useful information. Differential correlation of gene pairs is another mechanism by which the transcriptome is altered in disease.^12^ Deconvolution of the transcriptome to identify presence of various immune infiltrates and their contribution to disease processes is another insightful use of genome wide measurement of gene expression.^13^ These different approaches in studying the transcriptome provide complementary information and are critical both in establishing various aspects of disease pathogenesis and finding candidate drugs for therapy.

Several differential gene expression and a few gene co-expression studies in IgAN have been done,^14–17^ but a holistic evaluation of the transcriptome, involving differential gene expression, differential gene correlation, alteration of gene co-expression networks, and differential immune cell enrichment based on deconvolution of the transcriptome, is lacking. We therefore comprehensively evaluate the transcriptome of IgAN to identify different levels of dysregulation of the transcriptome and gain insights into disease mechanisms amenable to therapeutic intervention.

## Methods

### Data Acquisition and Preprocessing

Publicly available gene expression data from glomeruli of 22 healthy controls and 20 IgAN patients, used in PMID: 28646076 was used for our study.^18^ The raw affymetrix microarray files accessioned as GSE93798 were downloaded from the GEO database using GEOquery (version 2.54.1).^19^ The data files were read using the affy package (version 1.64.0)^20^ and quality control checks were done with the simpleaffy R package (version 2.62.0).^21^ We used probe level metrics, relative log expression and normalized unscaled standard error, and array level RNA degradation to assess quality of arrays. Arrays failing all 3 quality checks were to be removed. None of the samples failed the quality criteria.

The raw data was read with the simpleaffy, normalized and log transformed in batches using gene-chip robust multiarray averaging (GCRMA) and probe level data summarized to gene level expression with custom CDF for affymetrix HG-U133_Plus_2 arrays based on Entrez genes (hgu133ahsentrezgcdf_24.0.0). The normalized expression files were then merged. Since the microarray data was processed at 5 different time points, we checked for presence of batch effects using principal component analysis (PCA) and batch effects were corrected by empirical Bayes with the ComBat function of the sva package (version 3.34.0).^22^ PCA afterwards showed correction of batch effect but with one outlier IgAN sample, which was removed from subsequent analysis.

### Global Changes in the Transcriptome

A sample correlation heatmap with hierarchical clustering of samples based on disease status, and principal component analysis were used to identify global changes in the transcriptome in IgAN. The factoextra and Factominer R packages were used to extract genes most correlated with the first 2 principal components (PCs).^23,24^ An enrichment map was built to functionally analyze and visualize biologic processes and Reactome pathways enriched among genes with absolute correlation with PC1 > 0.5 and adjusted p-value of correlation < 0.05, using the grofiler-cytoscape-enrichment map pipeline as described.^25–27^

### Estimation of Immune Constitution

To characterize the immune composition of IgAN using the transcriptomic data, we used the imsig R package (version 1.0.0).^13^ In imsig, a network-based deconvolution using co-expressed gene modules associated with human immune cell populations is used to quantitatively estimate the relative abundance of seven immune cell populations (B cells, macrophages, monocytes, neutrophils, NK cells, plasma cells, and T-cells) and 3 biological processes (proliferation, translation and interferon response). A correlation threshold of 0.7 between imsig signature genes and our data was used for feature selection. Confirmation of the presence of the cell types was done by visualizing the network plot of cell types and processes, and evaluating the median correlation between imsig signature genes used for that cell type and genes in our dataset. Significance of the difference in means of the relative abundance of different cells and biological signatures was determined using a t-test with a p-value < 0.05 as criteria.

### Differential Gene Expression

Differential gene expression was performed with the Limma R package (version 3.42.2)^28^ using a Benjamini Hochberg adjusted p-value cutoff of 0.01 and a fold change of 2 as criteria for differentially expressed genes (DEG). The Cogena R package (version 1.22.0) was used to perform functional analysis of the differentially expressed genes.^29^ DEGs were grouped into co-expressed clusters and then enrichment analysis of the clusters was performed for KEGG pathway overrepresentation.

### Modular Co-expression Analysis

The CEMiTools R package (version 1.12.1)^30^ which uses several algorithms and other R packages for module detection, gene set enrichment of the modules in the groups being compared, and then functional analysis of the modules, was used for co-expression analysis. Briefly, unsupervised filtering of the genes using an inverse gamma distribution is first performed to select the most informative genes. Among the selected genes, a similarity criterion between gene pairs is determined using a soft thresholding power which is chosen based on the concept of Cauchy sequences. Dynamic tree cutting^31^ is then used to separate modules. Gene set enrichment analysis^32^ of identified modules is done with the fgsea R pakage^33^ to determine enrichment or repression of the identified modules in IgAN versus control patients. Functional analysis of the modules was then performed by assessment of overrepresentation using a hypergeometric test with the clusterprofiler R package^34^ for KEGG pathways (v7.01) downloaded from MSigDB.

### Network Analysis of Downregulated Metabolic Pathways in IgAN

To further demonstrate and provide granular details of the dysregulation of metabolic pathways in IgAN, we considered the metabolic pathways identified as aberrantly expressed in both functional analysis of DEG clusters and co-expression modules, and evaluated the preservation of internal structures of these pathways in IgAN compared with control. Each selected pathway was considered as a module, and the module in controls was used as a reference, and preservation of the module was assessed in IgAN using a composite network-based module preservation statistic called the Z-summary, with the module preservation function of the WGCNA R package (version 1.69).^35^ The Z-summary measures the preservation of the density and connectivity of genes in test network/module compared to the reference network.^36^ To visualize the internal structure of dysregulated pathways, we made pathway circle plots for the 2 least preserved pathways using a custom circle plot function available on https://horvath.genetics.ucla.edu/html/CoexpressionNetwork/ModulePreservation/Tutorials/ and described in Langfelder et al. ^36^ In this plot, nodes represent genes and edges indicate connection between genes. The size of the nodes is determined by intramodular connectivity of the gene, with larger sizes indicating higher intramodular connectivity. Genes with higher intramodular connectivity are more central in the pathway and behave as hub genes. The width of the edges represents the absolute value of the correlation between nodes, with red colored edges indicating positive correlation and blue colored ones for negatively correlated gene pairs.

### Differential Gene Correlation Analysis

We evaluated the IgAN transcriptome for ubiquity and type of differential gene correlation, and functional insights of differentially correlated genes. We filtered out genes with low variance by using the “filtergenes” function of the DGCA R package (version 1.0.2)^12^, keeping genes in the 80^th^ percentile of coefficient variation-a total of 4084 genes. Gene pair correlations among the 4084 genes in control was determined and used as reference for comparison for correlation of the same gene pairs in IgAN. The “ddcorall” function of DGCA determined gene pairs with altered correlation in IgAN compared with control. The function computes the difference in pairwise correlation in control versus IgAN, Z-transforms the difference and determines significance of the difference. We determined significance of differences in correlation after 100 permutations.

## Results

### Global Changes in the Transcriptome

Sample correlation heatmap and principal component analysis showed clustering of samples based on disease status (Fig 1a and Fig 1b). Genes contributing the most to PC1 include *ARHGAP33* and *DACT3-AS1* while *LOC105376856* and *NKAIN1* contributed the most to PC2 (Fig 1c and Fig 1d). Enrichment analysis showed that the most varying genes in the dataset, ie genes most significantly correlated with PC1, were enriched in cell adhesion, leucocyte differentiation, extracellular matrix organization and membrane transport (Fig 1e).

**Figure 1a:**
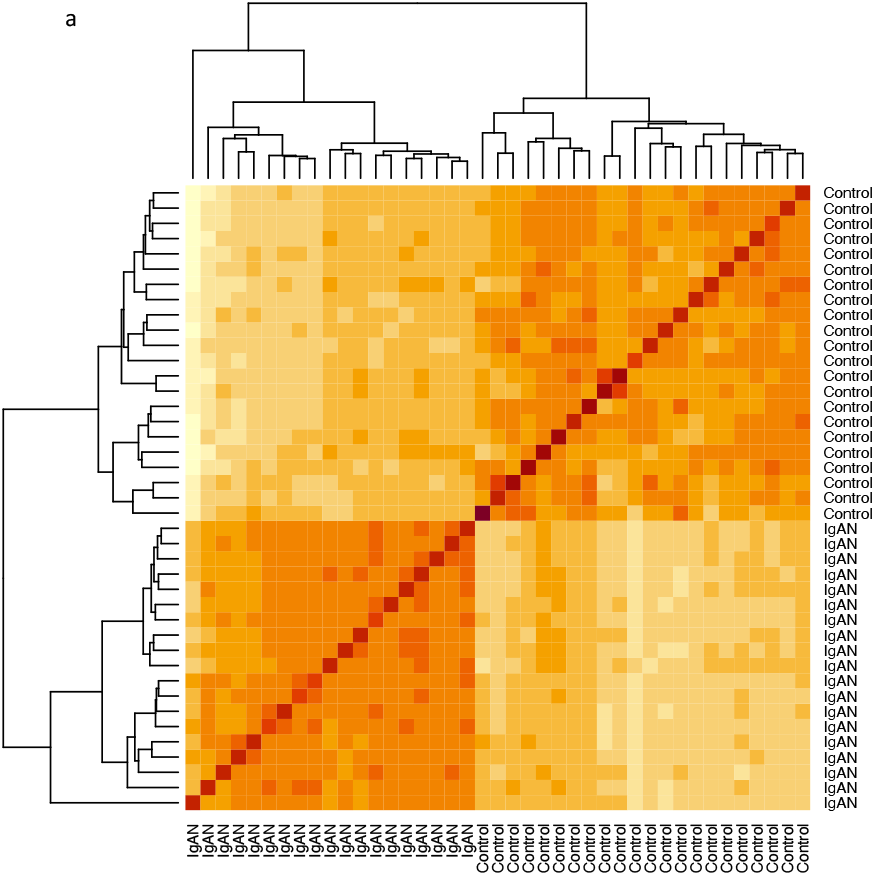
Sample correlation heatmap showing separation of samples by disease status.

**Figure 1b:**
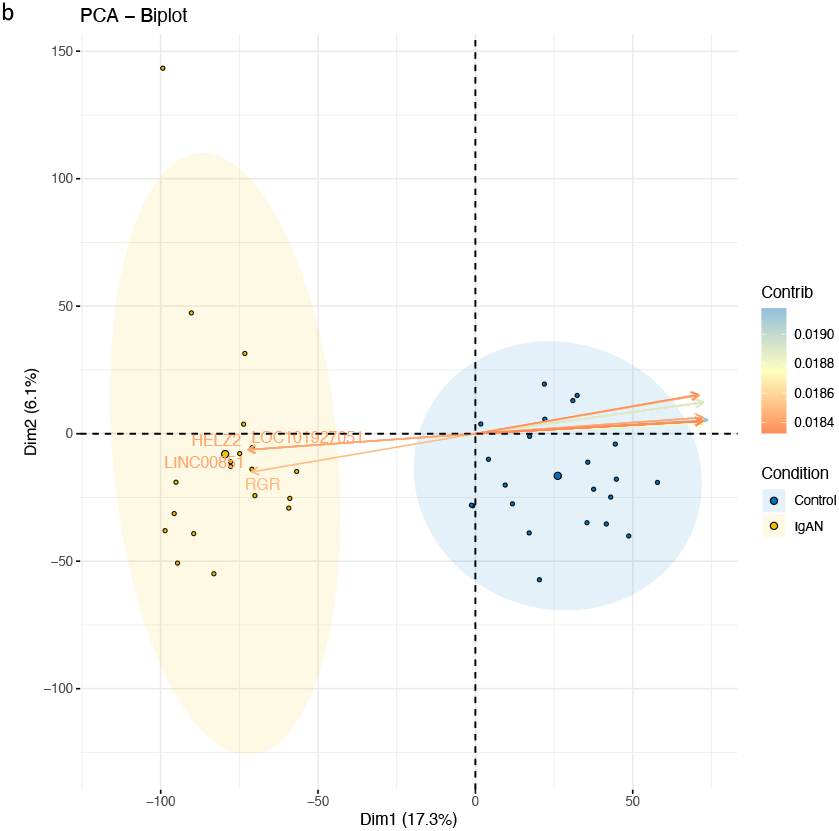
Biplot of 1^st^ 2 Principal Components with samples clustering by disease status, and genes(variables) colored by contribution to PC1 or PC2.

**Figure 1c:**
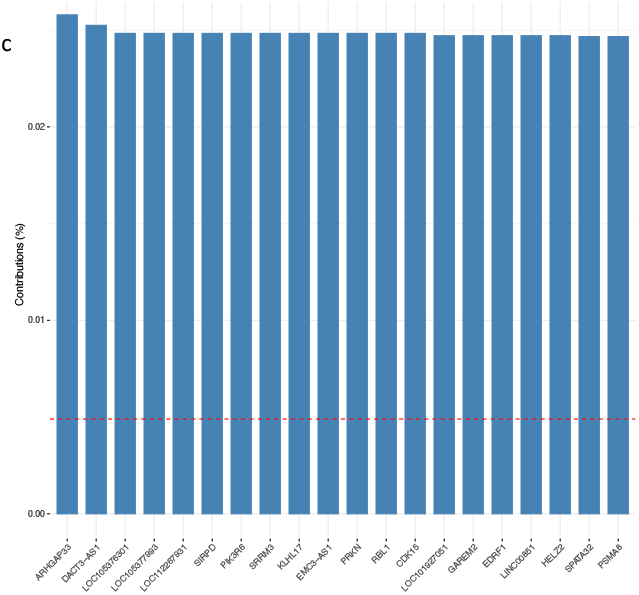
Variables with the highest contributions to the first principal component.

**Figure 1d:**
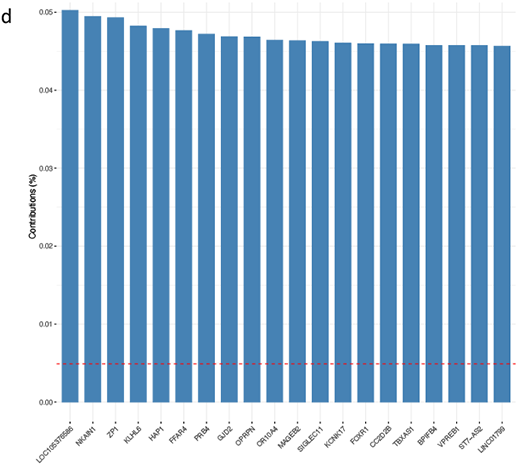
Variables with the highest contributions to the second principal component.

**Figure 1e:**
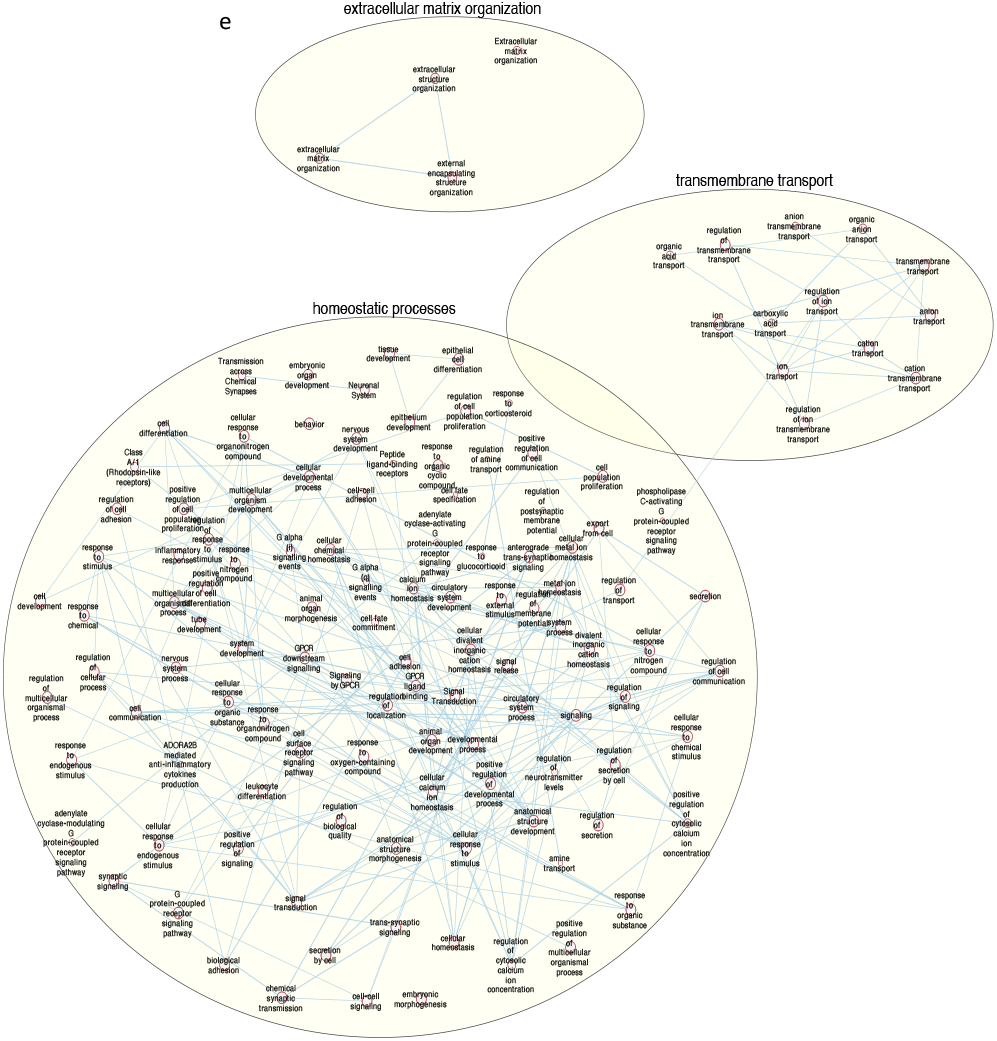
Enrichment map of biological processes and Reactome pathways of variables significantly correlated with the first principal component.

### Estimation of Immune Constitution

Immune environment analysis showed enrichment of B cells, macrophages, monocytes, plasma cells and T cells in IgAN (Fig 2a). Neutrophils and NK cells were excluded from further analysis because of their low median correlation and/or low abundance from the network plot (Fig 2A), as suggested by the authors of the package. The fold changes of enrichment of the estimated immune cells in IgAN were however not drastic, ranging from 0.07 for B-cells to 0.30 for plasma cells. The cell type with the highest average relative abundance in IgAN were monocytes. Additionally, interferon, translation and proliferation signatures were enriched in IgAN, with the proliferation signature showing the highest fold change compared with controls (Fig 2B).

**Fig 2a:**
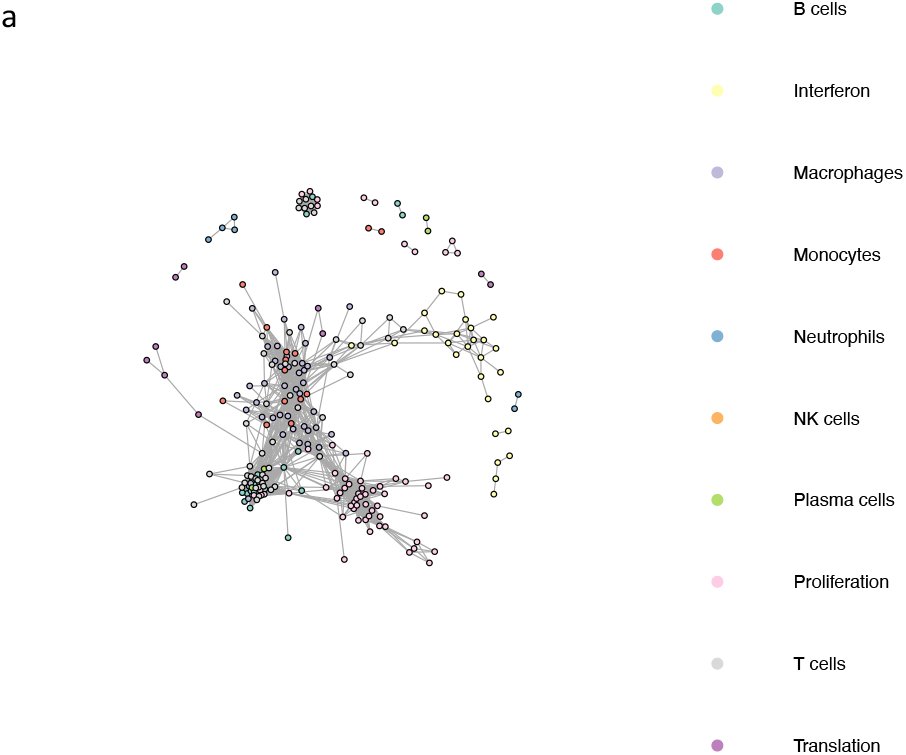
Network plot of various cell types and biologic processes in controls and IgAN.

**Fig 2b:**
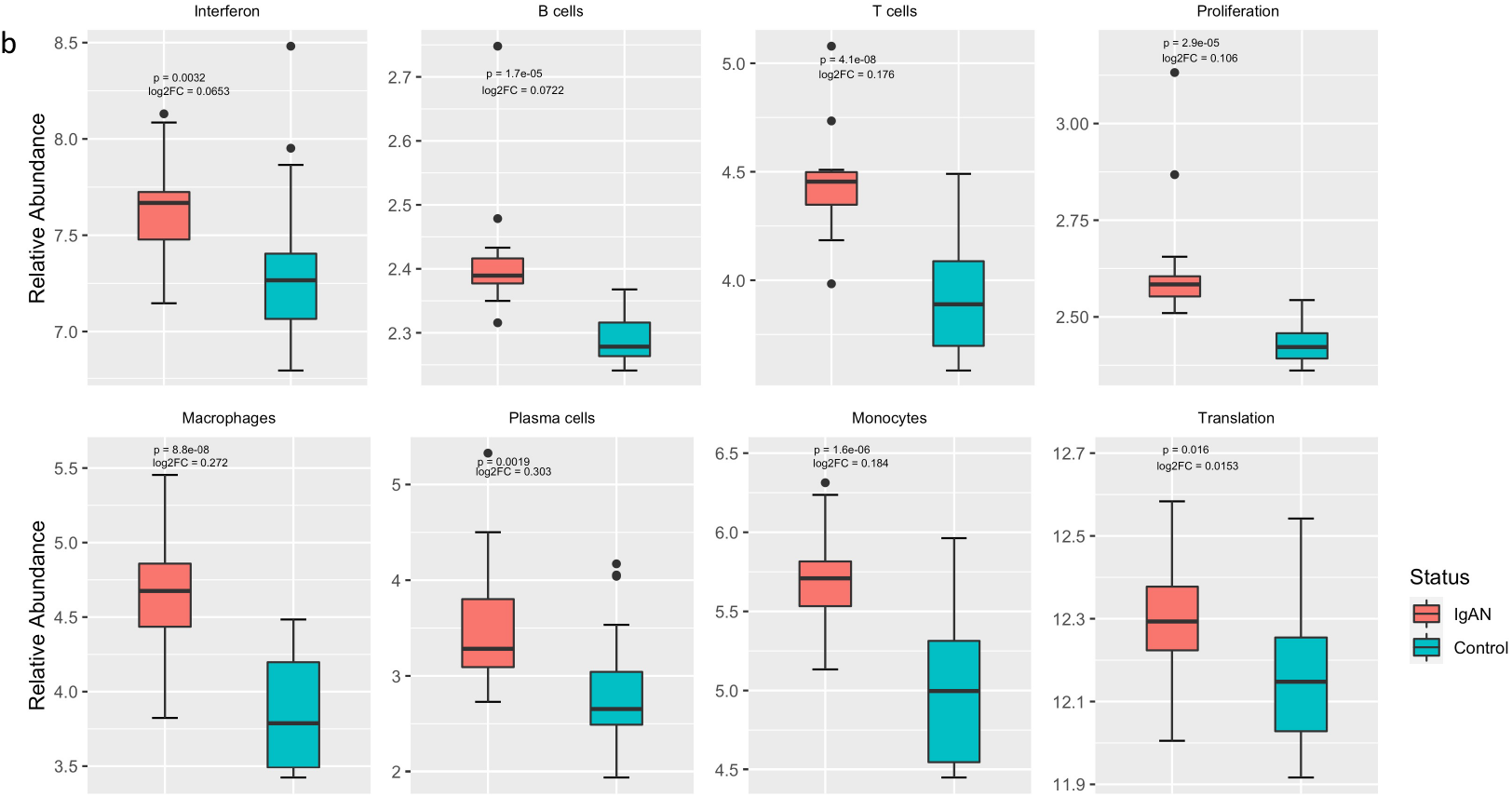
Differential abundance of immune cells and signatures for biologic processes in control versus IgAN.

### Differential Gene Expression

We identified 523 DEGs (Fig 3a and 3b). Of these, 174 were upregulated in IgAN and 349 were downregulated in IgAN. Upregulated genes included *AGTR1, ECM1, COL15A1, CXCl11, C1QA* and *C1QB* (Fig 3c). *PHGDH, FABP1, NFKBIA, PSAT1, BHMT* and *BHMT2* were among the downregulated genes (Fig 3d). The DEGs were clustered into 7 groups using partitioning around the medoid algorithm (Fig 3e). Three of the clusters consisted of upregulated genes and the other four consisted of downregulated genes. KEGG pathway enrichment was then performed for each cluster. Functional analysis revealed upregulation of known pathways involved in IgAN including focal adhesion, complement and coagulation cascades, hematopoietic lineage and extracellular matrix receptor interaction (Fig 3f). Cluster 6, a metabolic cluster, is a group of one hundred and ninety-one (191) co-downregulated genes showing enrichment of multiple amino acid metabolic pathways such as tryptophan metabolism, glycine, serine and threonine metabolism and PPAR signaling pathway (Fig 3f).

**Fig 3a:**
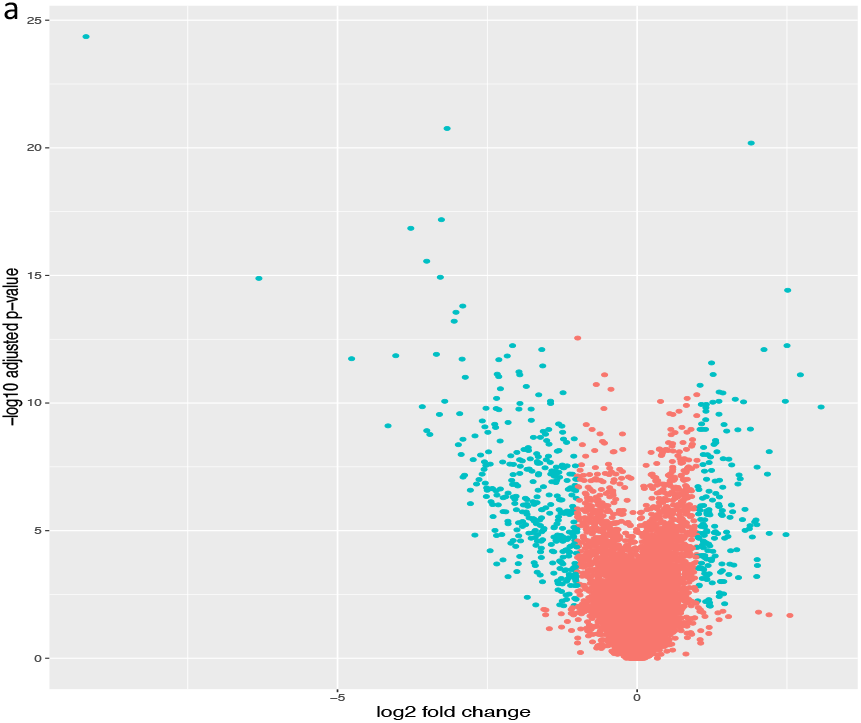
Volcano plot for differentially expressed genes using adjusted p-value < 0.01 and absolute fold change > 2, as criteria.

**Fig 3b:**
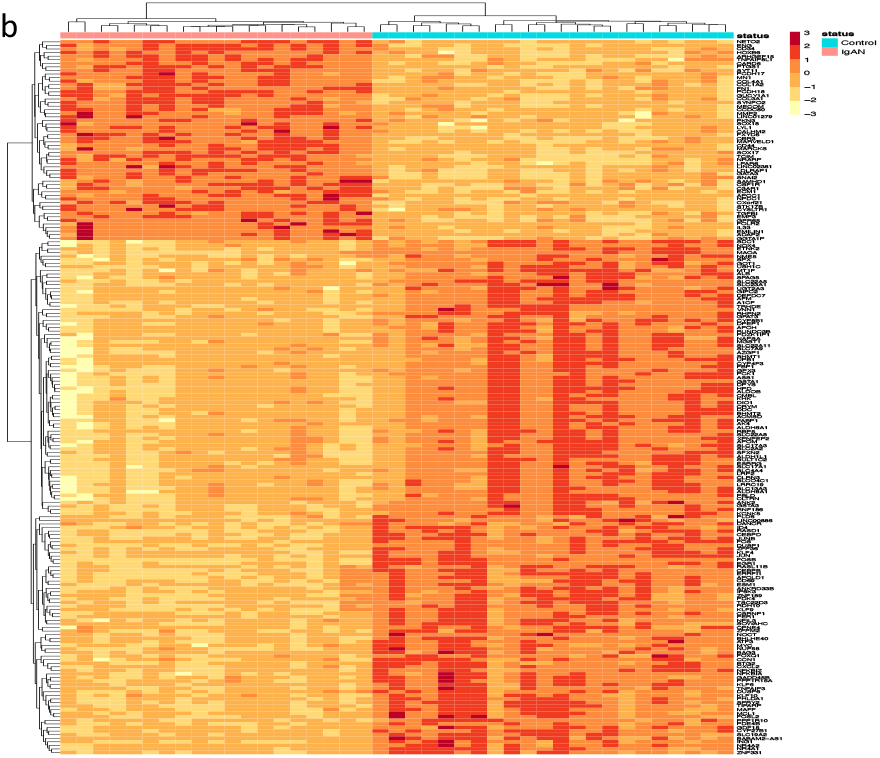
Heatmap of differentially expressed genes.

**Fig 3c:**
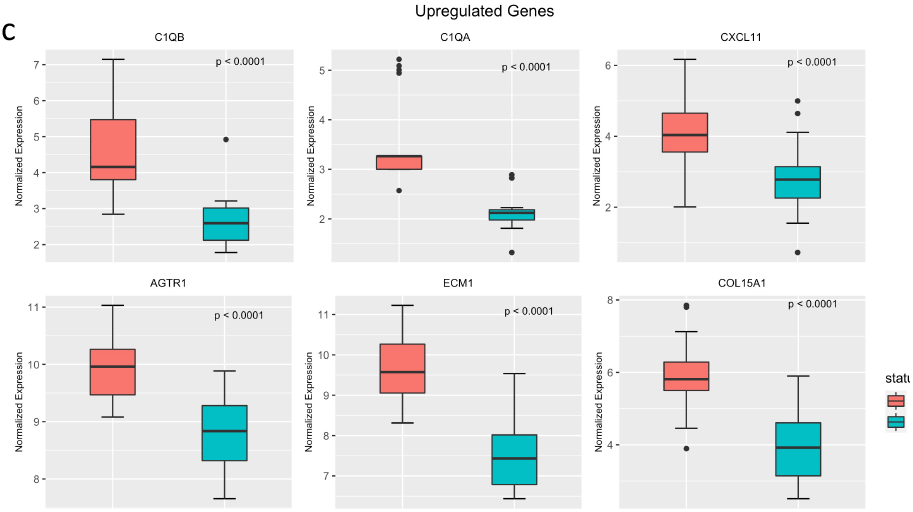
Select upregulated genes in IgAN.

**Fig 3d:**
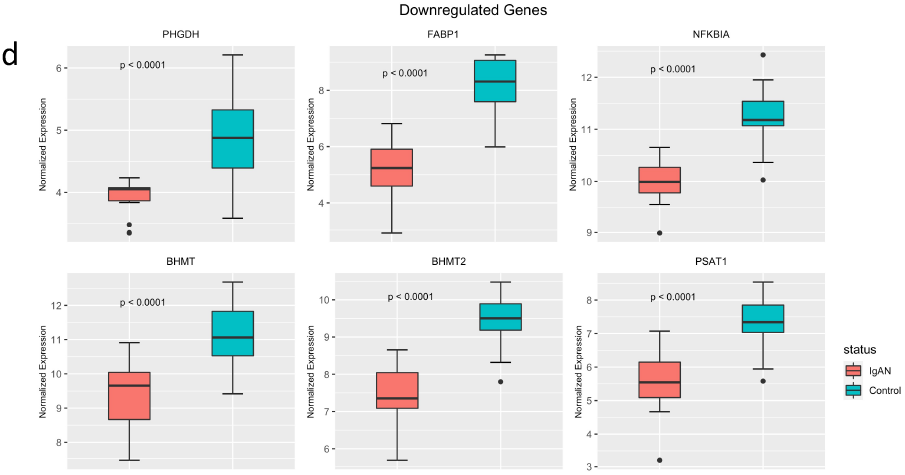
Select downregulated genes in IgAN.

**Fig 3e:**
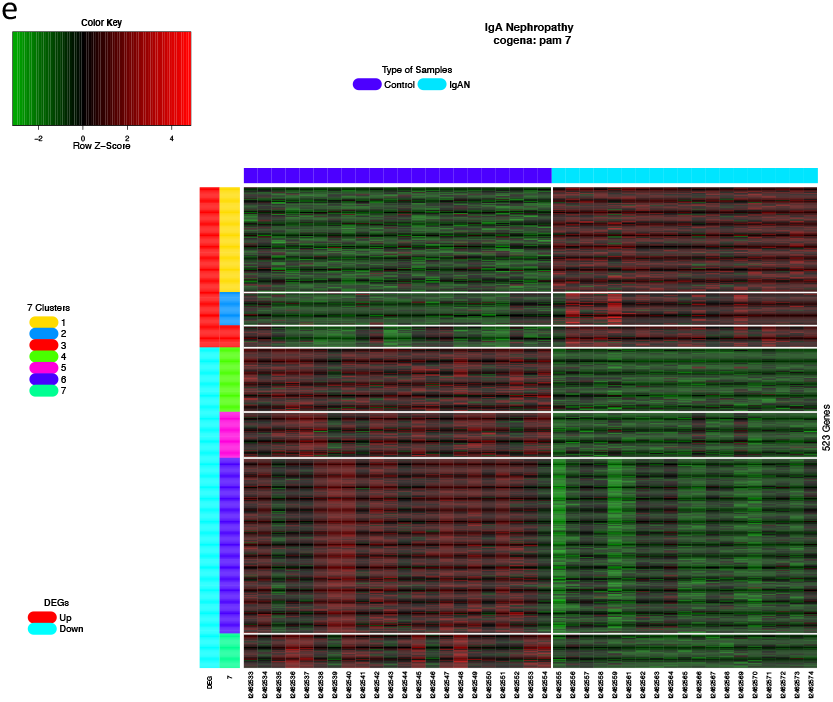
Seven clusters of co-expressed genes defined by partitioning around the medoids. Clusters 1-3 are upregulated in IgAN and clusters 4-7 are down-regulated.

**Fig 3f:**
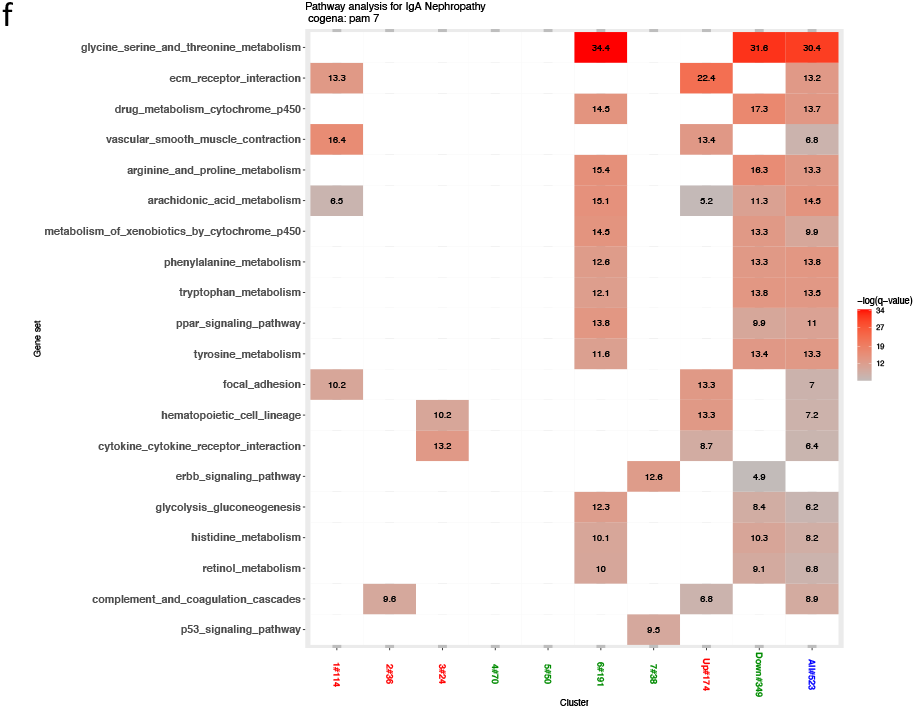
Functional analysis of enriched KEGG pathways in the seven co-expressed clusters.

### Modular Co-expression Analysis

After gene filtering, 1703 genes were selected for module identification. Six (6) modules of co-expressed genes were identified. Gene set enrichment analysis revealed significant enrichment of modules 2, 5 and 6 in IgAN, and significant repression of modules 1, 3 and 4 (Fig 4a). Functional analysis showed significant enrichment of KEGG pathways in modules 1, 2, 6 while modules 3, 4, 5, had no pathway enriched to statistical significance (Fig 4b).

**Fig 4a:**
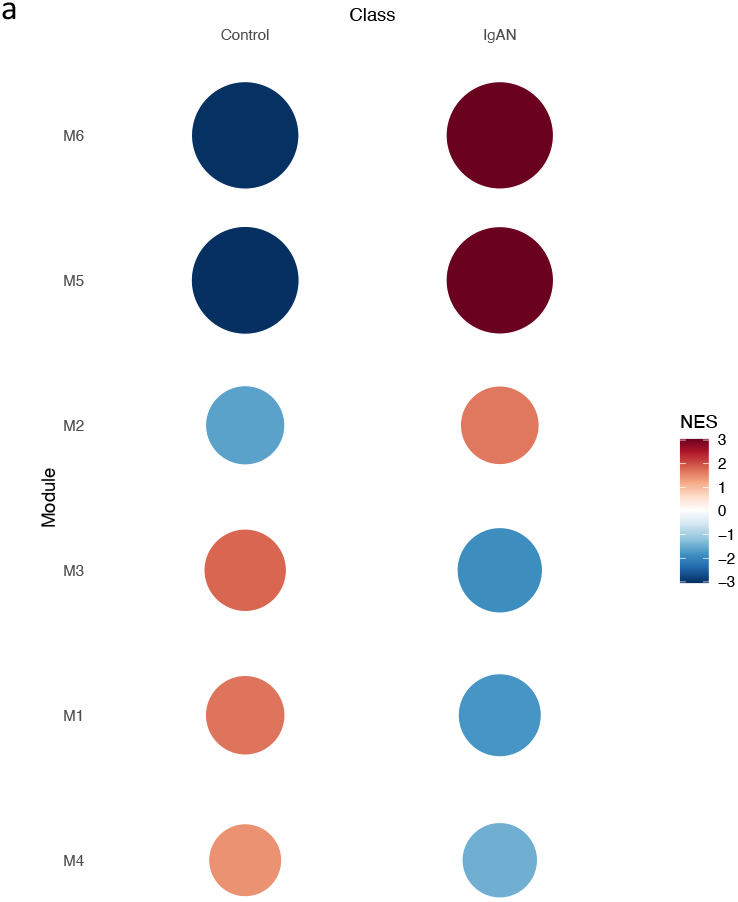
Gene set enrichment plots for 6 identified co-expressed modules, in controls and IgAN.

**Fig 4b:**
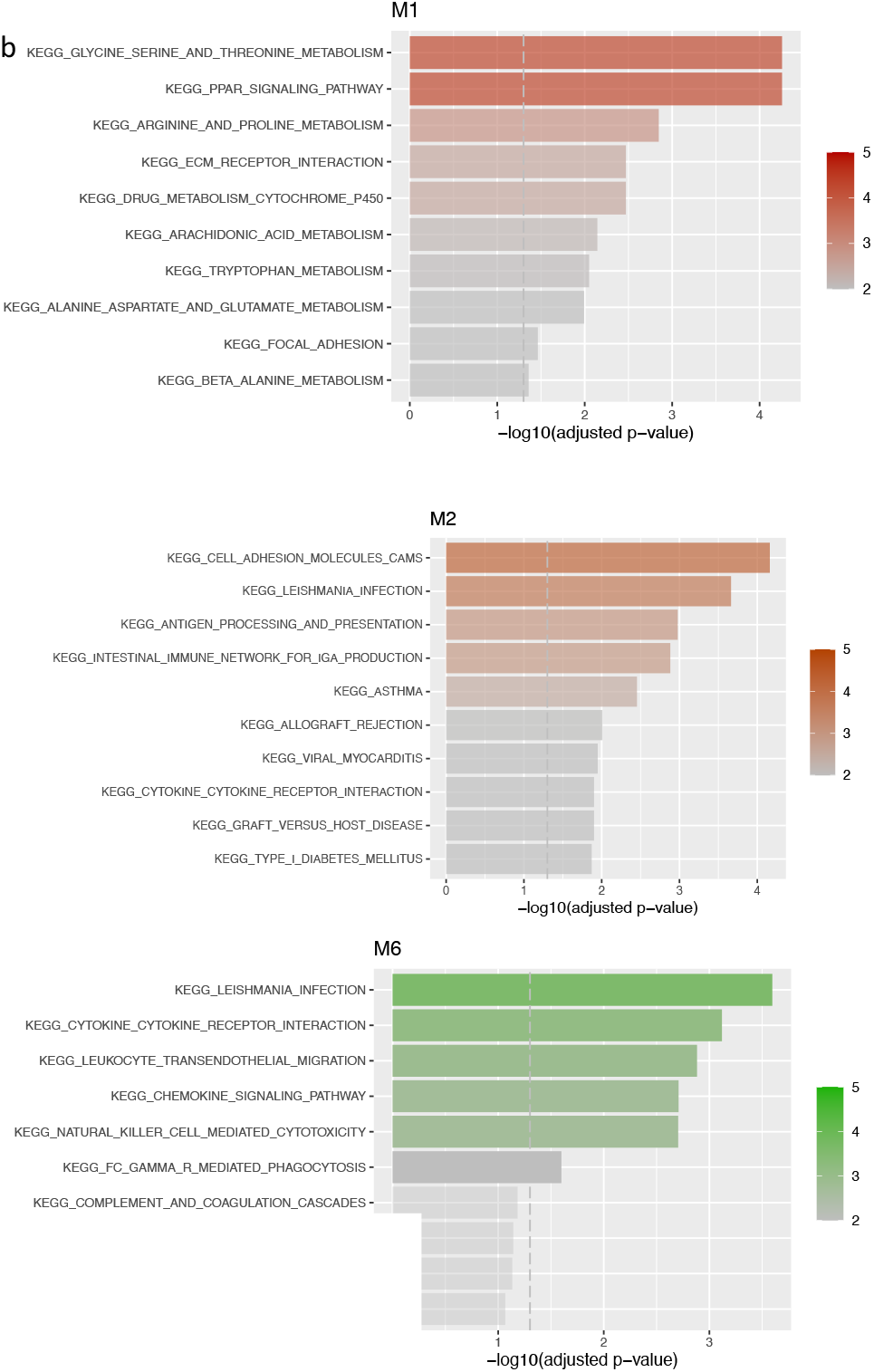
Enriched pathways in co-expressed modules 1,2,6 (from top to bottom).

**Fig 4c:**
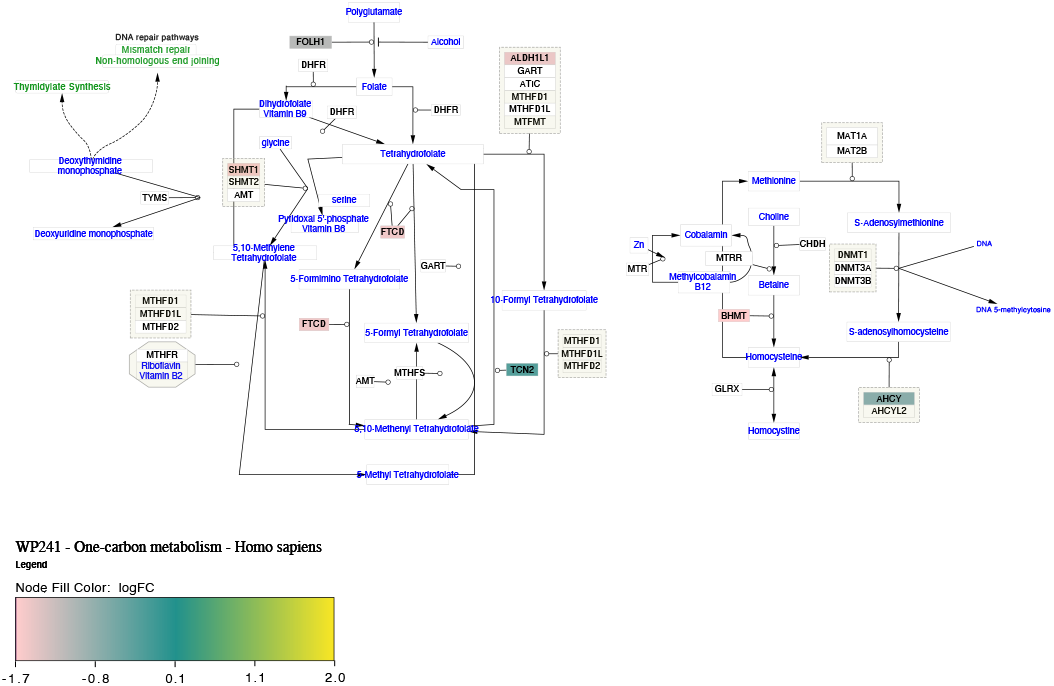
Network of one-carbon metabolism pathway. Module 1 genes in the pathway are colored by the log fold change of expression on differential gene expression.

Module one (1), a metabolic module, was repressed in IgA and showed functional enrichment enriched in amino acid metabolism such as glycine-serine-threonine metabolism, tryptophan metabolism and PPAR signaling. Since most of the amino acids involved in module 1 are involved in one-carbon metabolism, we imported the one-carbon metabolism network from NDEX using cytoscape. Module 1 genes were superimposed on the network as nodes and nodes colored by fold chain (Fig 4b), showing predicted repression of the pathway. Modules 2 and 6 were immune modules, enriched in IgAN and showed functional enrichment in expected immune processes such as leucocyte migration, cytokine-cytokine receptor interaction, complement activation and coagulation cascade, NK-cell mediated cytotoxicity, chemokine signaling and FC-gamma receptor mediated phagocytosis. Curiously, in module 2, we identified enrichment of the intestinal network for IgA production in IgAN.

### Network Analysis of Downregulated Metabolic Pathways

We selected glycine, serine and threonine metabolic pathway, PPAR signaling pathway, arginine and proline metabolic pathway and tryptophan metabolism, as pathways enriched in both the repressed metabolic module and downregulated DEG metabolic cluster in IgAN, for analysis of preservation. All the pathways showed weak to moderate evidence of preservation (2 < Z-summary <10) (Fig 5a). PPAR signaling and glycine, serine and threonine metabolic pathways were the least preserved. Circle plot of the pathways revealed some interesting findings. There is a loss of intramodular connectivity (hubness) of *PPARA* as well as a reduction in the correlation between transcript levels between PPARA and its receptor RXRA in IgAN (Fig 5b). *PPARA* however seems to gain correlation with *RXRB* in IgAN. On the other hand, *PPARG*, gains intramodular connectivity in IgAN and shows an increase in its correlation with its receptor *RXRG* (Fig 5b).

In controls, transcript levels of phosphoglycerate dehydrogenase (*PHGDH)*, which is the rate limiting step in serine synthesis, correlates with *SHMT1* which reversibly converts serine to glycine in the cytosol (Fig 5c). This correlation appears lost in IgAN. Similarly, in IgAN, *SHMT2*, the mitochondrial form of serine hydroxymethyltransferase, thought to be the primary source of intracellular glycine, shows loss of positive correlations with its cytosolic isoform *SHMT1* and with *PSAT1*, an upstream protein in serine synthesis whose products feeds into the *SHMT2* reaction (Fig 5c).

**Fig 5a:**
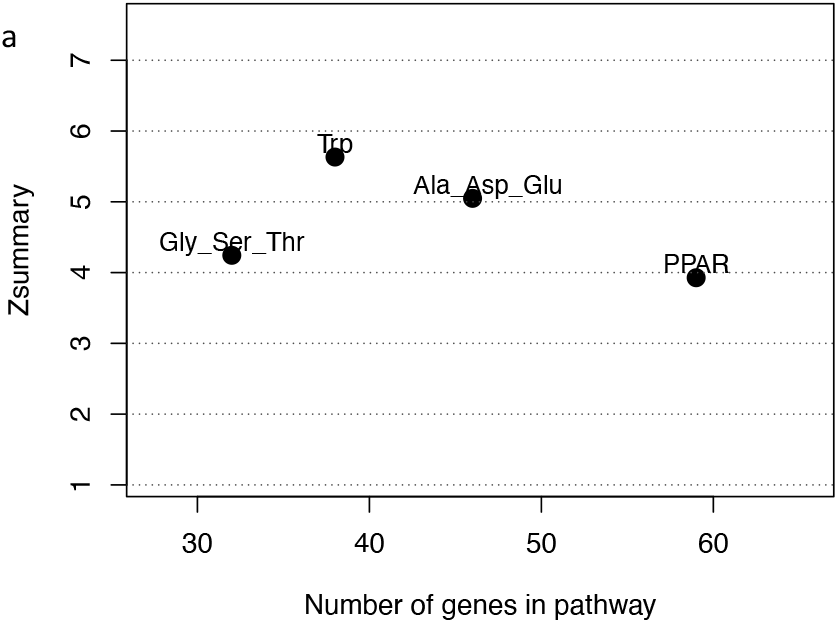
Composite preservation statistic, Zsummary, for KEGG pathways repressed on both modular co-expression and differential gene expression analyses, all showing weak to moderate preservation (2 < Zsummary < 10). Gly_Ser_Thr: Glycine, serine, threonine metabolism. Trp: Tryptophan metabolism. Ala_Asp_Glu: Alanine, aspartate, glutamate metabolism. PPAR: Peroxisome proliferator-activated receptor signaling.

**Fig 5b:**
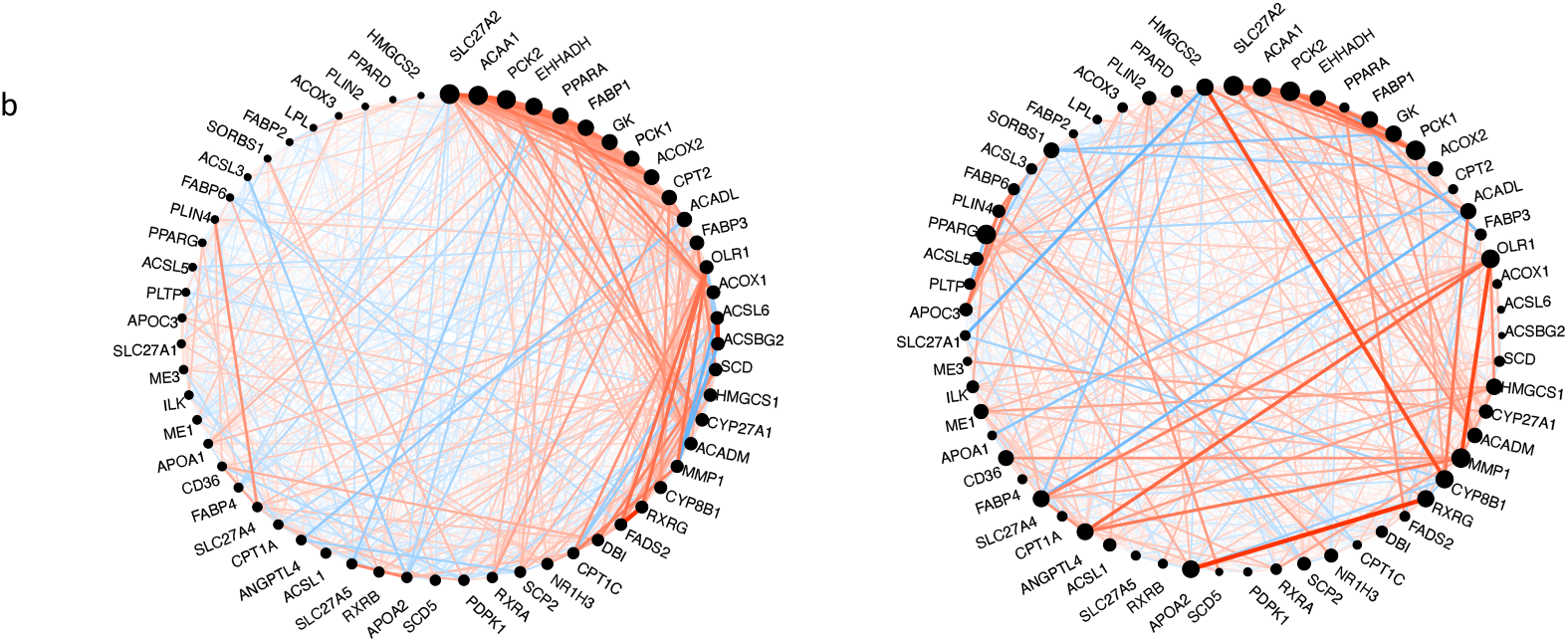
Circle plot of PPAR-signaling pathway using control (left circle) as reference and showing network alteration in IgAN (right circle). Nodes represent genes and node size is proportional with intramodular connectivity (hubness). Edges represent connection between genes. Red edges indicate positive correlation, and blue edges indicate negative correlation. Edge thickness represents absolute correlation. Network alterations in IgAN include loss and gain of hubness in *PPARA* and *PPARD* respectively.

**Fig 5c:**
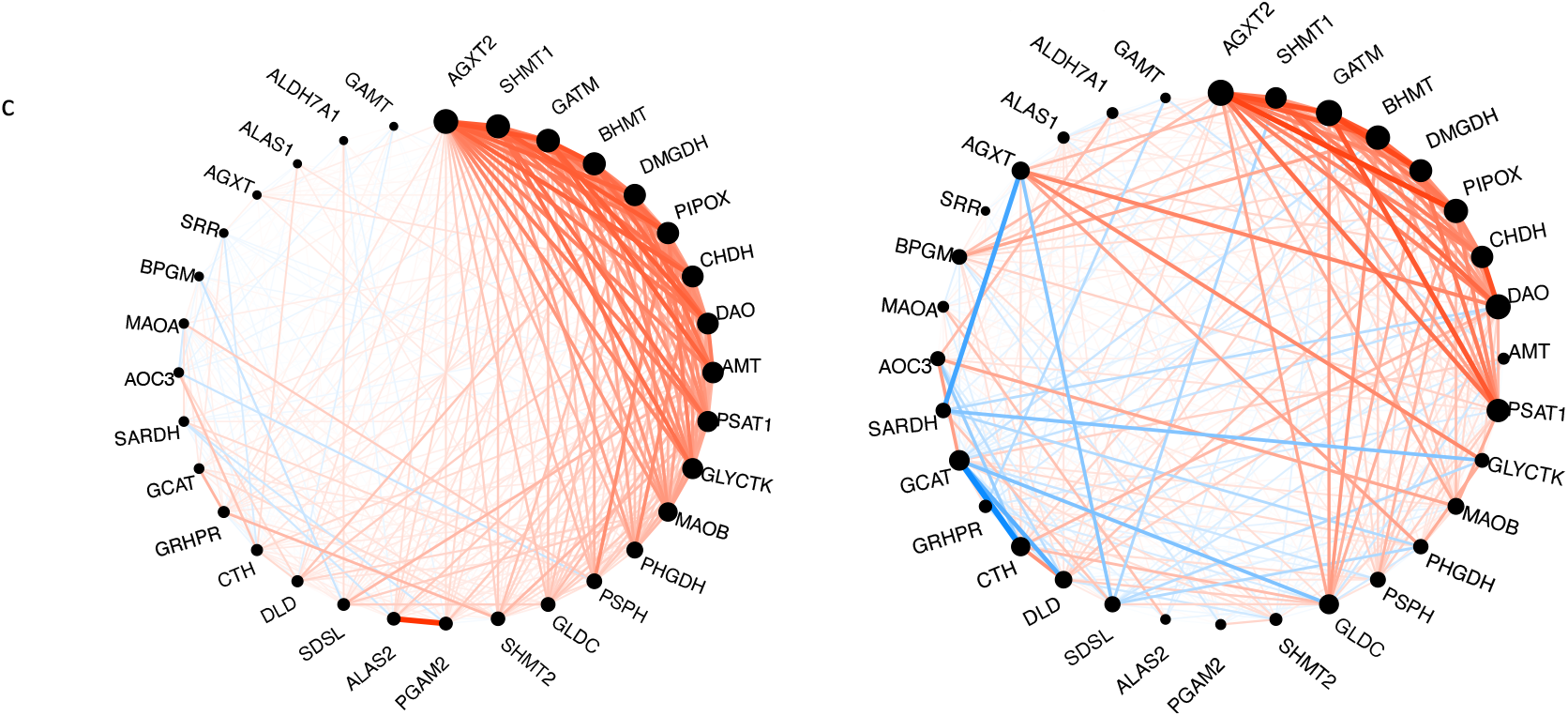
Circle plot of glycine-serine-threonine pathway using control (left circle) as reference, showing network alterations in IgAN (right circle) including loss of correlation between *PHGDH*, the rate determining step in serine synthesis, and *SHMT1*, which converts serine to glycine.

### Differential Gene Correlation Analysis

There were 1560 gene pairs with significantly altered correlations in IgAN, indicating that differential gene correlation is a common phenomenon in IgAN (Fig 6a). The different classes of pairwise alteration of correlation are shown in Fig 6b. We find that several of the significantly different alterations involve matrix producing and matrix degrading genes. A notable example is the correlation between the matrix producing, COL14A1 and matrix degrading, MMP1, which is a strong positive correlation in controls, but the correlation is lost in IgAN (Fig 6c). A similar loss of positive correlation between MFAP2 which is matrix producing and MMP1 in IgAN.

**Fig 6a:**
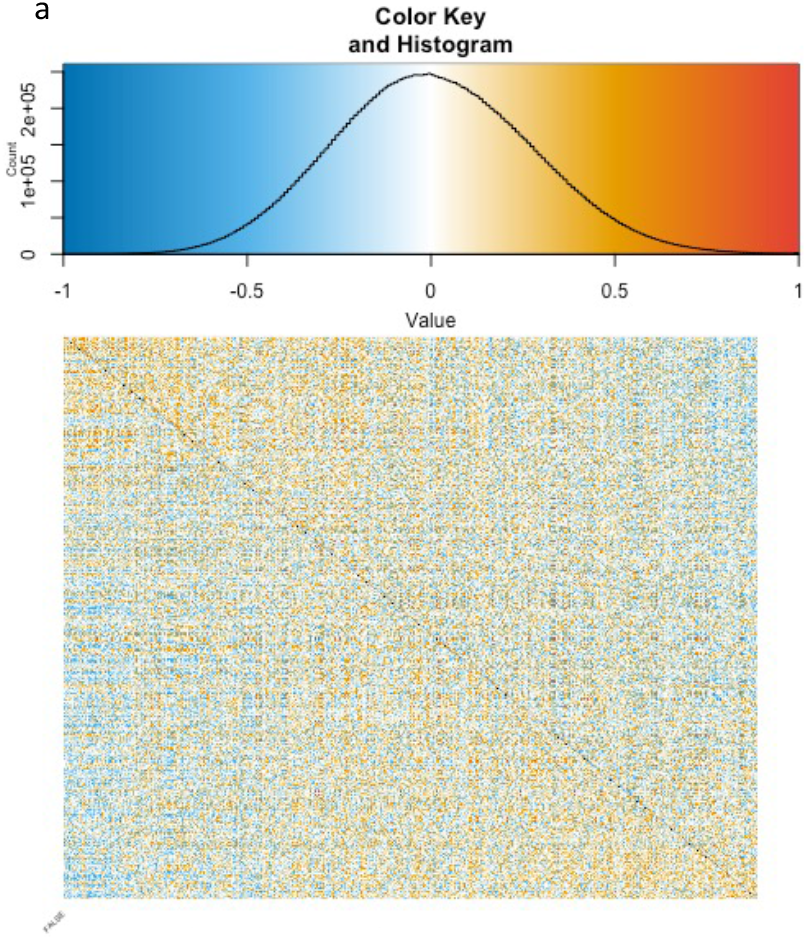
Correlation heatmap of gene pairs. Lower triangle is for controls and upper triangle is for IgAN.

**Fig 6b:**
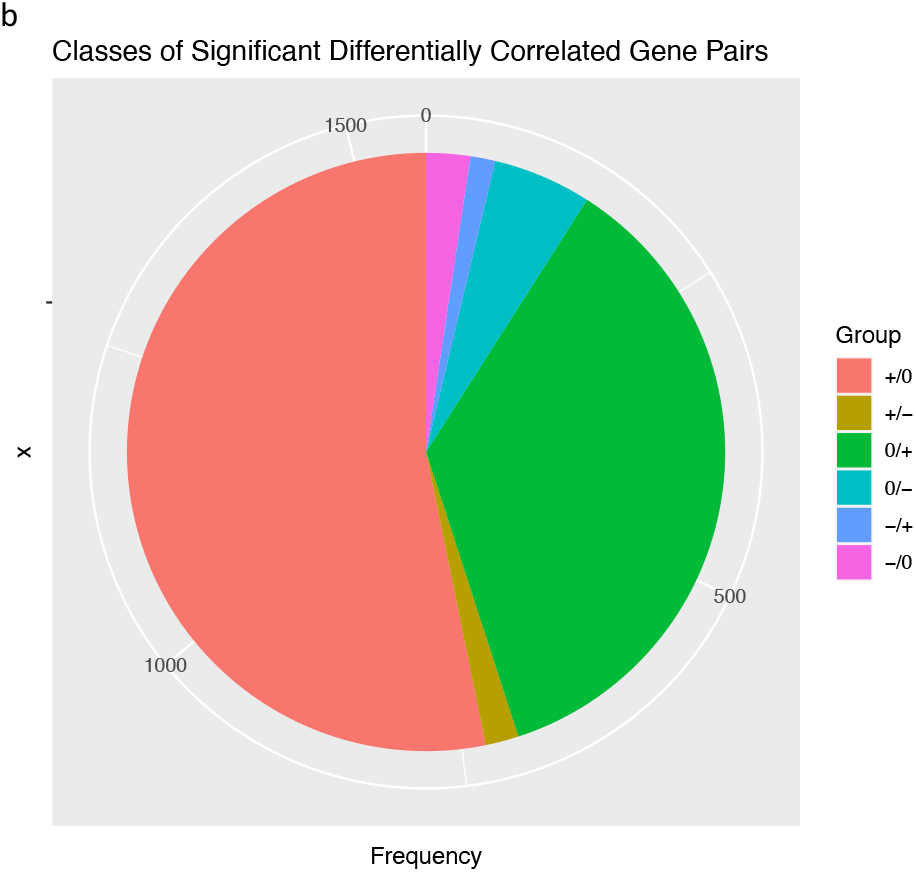
Counts of classes of differentially correlated genes +/0: Positively correlated pairs in control, no correlation in IgAN. +/- : Positively correlated pairs in control, negative correlation in IgAN. 0/+ : No correlation in control, positively correlated pairs in IgAN. 0/- : No correlation in control, negatively correlated pairs in IgAN. -/+ : Negatively correlated in controls, positively correlated in IgAN. -/0 : Negatively correlated pairs in control, no correlation in IgAN.

**Fig 6c:**
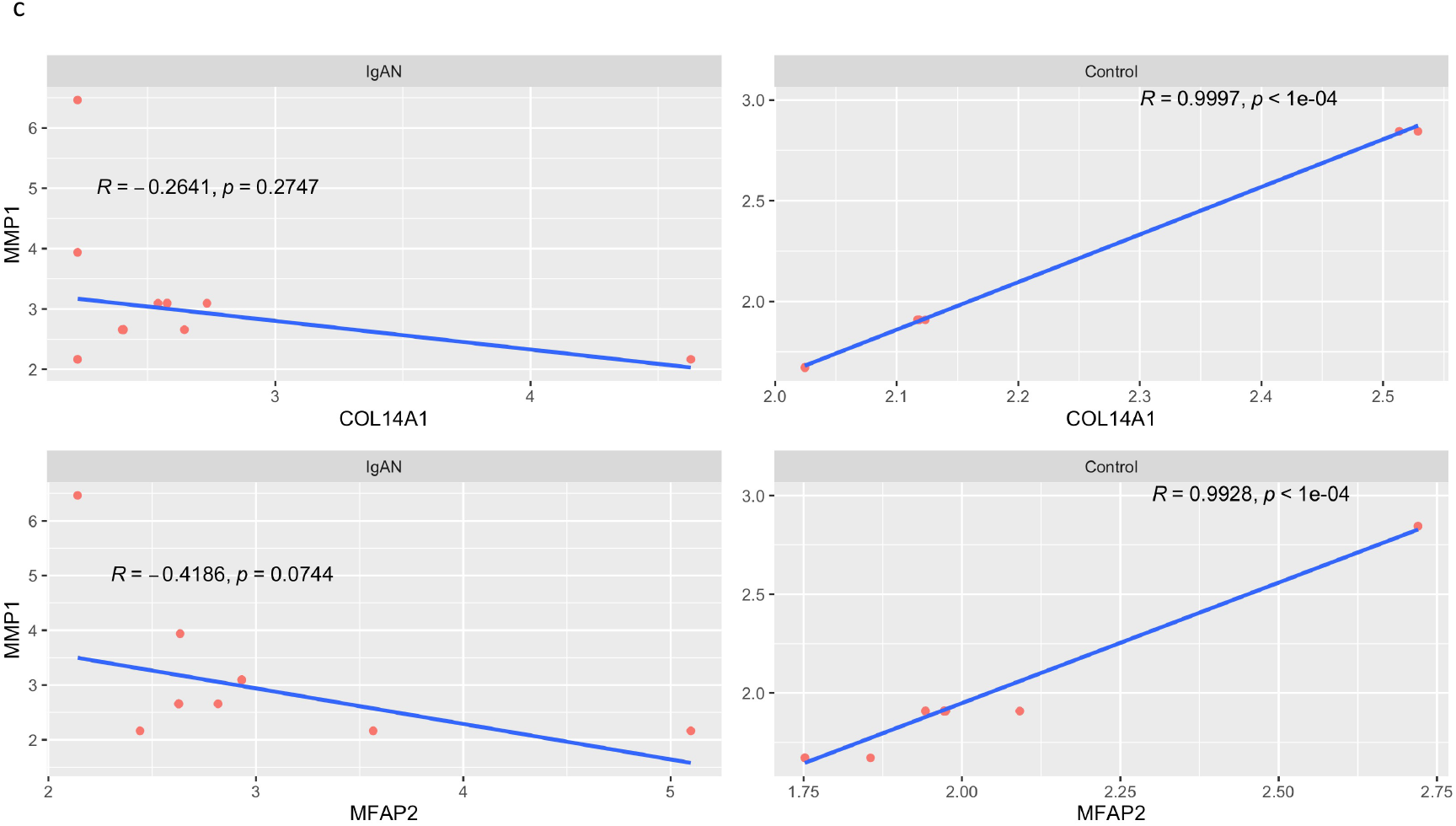
Positive correlation between COL14A1 and MMP1 in control (top right) and loss of the correlation in IgAN (top left). Similar loss of correlation seen between MMP1 and MFAP2 in IgAN.

## Discussion

IgA Nephropathy is the most common type of glomerulonephritis and currently considered an autoimmune disease with a four-hit hypothesis elaborating the mechanism of the disease. It is a progressive disease with no definitive cure. We have comprehensively characterized different ways the transcriptome of IgAN deviate from normal, with the aim of identifying features targetable by therapy.

Traditional immunosuppressive and B-cell depleting therapies have not shown significant efficacy in IgAN, and instead cause considerable adverse effects in patients.^37–39^ On estimation of the immune constitution of IgAN, we find significantly higher abundance of inflammatory cells and strength of signatures for translation, proliferation and interferon signaling. However, the fold changes for the inflammatory cells and biological signatures were considerably low. While this might be a feature of just this dataset, especially since the immune infiltrate of IgAN typically varies on histology of renal biopsies, there is the possibility that the low fold changes of the biological processes partly explain why immunosuppressive agents, which target different points along these signatures, have not shown convincing efficacy in IgAN. Conversely, considering the ranges of the strength of these signatures in our analysis, it may be useful to estimate abundance of inflammatory cells and strength of biological signatures as biomarkers to stratify patients in future clinical trials for IgAN therapy. Monocytes were the most abundant in IgAN while plasma cells had the highest fold change compared to controls. The latter finding is interesting considering that plasmablasts, the precursor cells of plasma cells, were found to be elevated in serum of IgAN patients, a finding distinct from other glomerulonephritis.^40^ Circulating plasmablasts have been found to correlate with level of proteinuria in IgAN patients^41^ and these findings suggests possible utility of plasma cell targeting therapies in IgAN. Mesangial IgA is considered to be derived from circulation, evidence of which is supported by clearance of abnormal IgA in allografts from donors with subclinical IgA nephropathy few weeks after transplant.^42–44^ Our immune module 2, which is upregulated in IgAN, shows enrichment of the intestinal network for IgA production. While this pathway has been known to be enriched in genome wide association studies of IgAN patients^45–47^ and functional analysis of transcriptome of blood derived cells^48^, transcriptomic evidence of its enrichment in glomerular biopsies is curious. Renal infiltrates of CD19+CD5+ B cells, which secrete IgA and inflammatory cytokines, have been described in IgAN patients.^49^ This finding suggests, that there might be need for renal suppression of IgA production in addition to current efforts designed to treat IgAN by suppressing intestinal IgA production with long-acting steroids.^50^ We find differential expression of immune modulatory genes and enrichment of inflammatory pathways in line with the autoimmune nature of the disease. Complement and coagulation cascade, chemokine signaling, cytokine-cytokine interaction and leucocyte transendothelial migration, already established features of the disease, were all enriched in our analysis. Among upregulated genes were *AGTR1* and *CXCL11*. Failure to suppress *AGTR1* expression after prolonged exposure to Gd-IgA1 has been associated with increased inflammatory and proliferative processes in mesangial cells.^1^ NFKBIA which we found to be downregulated in IgAN, encodes IkBa, a potent inhibitor of NF-kB, a key driver of inflammation.^51^ Targeted therapy of these gene products may be beneficial in modulating the immune activity in IgAN. Mesangial sclerosis is a feature of IgAN and results from increased matrix synthesis in activated mesangial cells and is an adverse prognostic indicator.^52,53^ Not surprisingly, there was enrichment of extracellular matrix organization among the genes contributing to the most variation in the global transcriptome of IgAN. Additionally, focal adhesion was seen in the cluster 1 of upregulated genes and immune module 1 enriched in IgAN. We found increased expression of basement membrane and extracellular matrix proteins *ECM1* and *COL15A1* in IgAN in agreement with proteomic studies evaluating the nature of the matrix in IgAN.^54^ We identified a difference in the correlation between expression of matrix-producing and matrix degrading genes in controls versus IgAN, on differential gene correlation analysis. While increasing expression of the extra-cellular matrix producing genes such as *COLA14* and *MFAP2*, is associated with an increase in expression of the matrix degrading gene *MMP1* in controls, this correlation is not observed in IgAN. This demonstrates the altered balance between matrix synthesis and degradation in IgAN. Identifying the drivers for this altered correlation may be useful in further understanding and targeting mesangial sclerosis in IgAN.

Metabolic reprogramming has been associated with the inflammatory phenotype in diseases such as systemic lupus erythematous, multiple sclerosis (MS), rheumatoid arthritis, psoriasis and some glomerulopathies, and has been the target of therapeutic interventions.^55^ For instance, modulation of reprogrammed metabolic pathways with dimethyl fumarate (DMF) in multiple sclerosis, psoriasis and SLE, switches the immune phenotype to an anti-inflammatory profile.^56^ In MS patients, modulation of lipid and fatty acid metabolism by DMF was associated with reduction in absolute lymphocyte counts and levels of subsets of cytotoxic T-cells.^57^ In autosomal dominant polycystic disease, alterations of glucose, fatty acid and amino acid metabolism are suspected to contribute to cyst proliferation and growth.^58^ Among our DEG clusters and co-expressed modules, cluster 6 and module 1, are a group of co-regulated and under expressed genes, and a repressed module of co-expressed genes, respectively, which show repression of amino acid metabolic pathways such as glycine-serine-threonine metabolism, alanine-aspartate-glutamate metabolism and tryptophan metabolism, in IgAN. Serine, glycine, threonine metabolism is involved in many pathways including the one-carbon cycle, which provides substrates for nucleotide synthesis, methylation reactions and anti-oxidant defense.^59^ Serine metabolism pathway genes such as *PSAT1* and *PHGDH* were among the downregulated genes in our analysis. PHGDH catalyzes the committed step in de novo serine synthesis, and is critical for nucleotide synthesis and one-carbon metabolism.^60^ Downregulation of *BHMT* and *BHMT2*, as seen in our analysis, may impair their critical roles of maintaining methionine levels in the cell, reducing levels of homocysteine, and regulating cellular volume and tonicity through control of the cellular osmolyte betaine.^61^

Using network analysis, we demonstrated with more granular evidence, the dysregulation of these metabolic pathways, in the form of loss of pathway preservation of in IgAN, as well as impairment of co-ordination of transcript levels of pathway genes. The relevance of the results of our analyses is underscored by the demonstrated efficacy in pre-clinical studies targeting these aberrant metabolic pathways in IgAN and other glomerulonephritides.

Gong et al described an injury pattern in rat mesangial cells induced by surfactants which cause proinflammatory and fibrotic transcriptomic changes akin to IgAN, as well as repression of amino acid metabolism. Curiously, pre-treatment of mesangial cells with amino acid supplementation, significantly reduced the surfactant induced inflammatory and fibrotic changes.^62^ The above suggests that rewiring these aberrant metabolic profiles in IgAN using small molecule inhibitors or nutritional interventions may be therapeutically beneficial in controlling the inflammatory phenotype and enhancing podocyte and tubulointerstitial cell survival.

Our results and discussion so far demonstrate immune cell and signature enrichment, altered balance in collagen synthesizing and degrading genes, and repression of amino acid metabolism in IgAN. The peroxisome proliferator activator pathway is linked to all these three aberrations and serves as a potential nodal point to control these aberrant processes.^63^ PPARs are nuclear receptors which regulate transcription of gene cassettes after ligand binding and heterodimerization with retinoid X receptors, mediating a coordinated response to a specific stimulus.^63^ There are three PPAR isoforms and together, they have anti-inflammatory effects, modulate lipid, glucose and amino acid metabolism, and exhibit anti-fibrotic activity.^64–67^ Not surprisingly, our analyses show that amino acid metabolic pathway genes and PPAR pathway genes cluster together in module 1, and are both repressed in IgAN. We also find loss of preservation of the PPAR pathway in IgAN, using network analysis, and illustrate aberrant transcriptional network connectivity between PPARs and their respective retinoid X receptors. This suggests a role for targeting PPAR activity in IgAN treatment. Indeed, several studies have shown efficacy of PPAR agonists in pre-clinical and clinical studies in IgAN. Rosiglitazone, a *PPARG* agonist potentiated the anti-inflammatory effect of angiotensin receptor blockers in proximal tubular epithelial cells activated by conditioned media from human mesangial cells incubated with IgA1 from IgAN patients.^68^ Administration of pemafibrate, a selective peroxisome proliferator-activated receptor-alpha modulator, to IgAN patients was associated with reduction in proteinuria.^69^ Of note, we see an increase in intramodular connectivity of *PPARD* in IgAN, which is a curious finding considering the variation in the biology of PPARs. Ligand binding results in PPAR dissociation from its co-repressors, and binding to co-activators, allowing binding of the complex to response elements on DNA. PPARD, unlike *PPARA* and *PPARG*, is able to bind DNA while still bound to co-repressors, in a ligand independent manner and exhibit competitive antagonism of ligand induced DNA binding by *PPARA* and *PPARG*.^70^ The relevance of increased connectivity of PPARD in the context of this known phenomenon and how it affects the effects of PPAR action in IgAN would have to be studied further. However, it may mean that PPAR targeted therapy may be more efficacious by stimulating multiple receptors to prevent this competition. Additionally, we find a reduction in expression of *FABP1*, which may have an effect on patient response to PPARA agonists. FABP1 controls lipid metabolism and loss of its expression reduces *PPARA* activation in response to agonist treatment.^71^

In summary, we have used complementary bioinformatic approaches to describe how the transcriptome of the glomeruli of IgA nephropathy deviates from normal glomeruli. Our approach demonstrates enrichment of immune processes such as complement and coagulation system, focal adhesion and cytokine receptor interactions, and enrichment of various immune cells and signatures, recapitulating known mechanisms of the disease. We identify transcriptomic evidence of enrichment of intestinal network for IgA synthesis in the glomeruli of IgAN patients. Our analysis describes alteration of the well-coordinated and correlated expression of matrix producing and matrix degrading genes. We additionally identify transcriptomic evidence of metabolic reprogramming in IgAN, predominantly involving repression of expression and dysregulation of networks of amino acid metabolic pathways together with the peroxisome proliferator activator receptor pathways. Our study highlights alterations in amino acid metabolism and PPAR pathways and suggests the use of drugs and other interventions targeting these alterations in treating IgAN.

